# SeqForge: A scalable platform for alignment-based searches, motif detection, and sequence curation across meta/genomic datasets

**DOI:** 10.1101/2025.08.12.669971

**Authors:** Elijah R. Bring Horvath, Jaclyn M. Winter

**Affiliations:** Department of Pharmacology and Toxicology, University of Utah, Salt Lake City, Utah, 84112, United States

**Keywords:** Genomics, bioinformatics, microbes, genome mining, BLAST, metagenomics

## Abstract

**Background:** The rapid increase in publicly available microbial and metagenomic data has created a growing demand for tools that can efficiently perform custom large-scale comparative searches and functional annotation. While BLAST+ remains the standard for sequence similarity searches, population-level studies often require custom scripting and manual curation of results, which can present barriers for many researchers.

**Results:** We developed SeqForge, a scalable, modular command-line toolkit that streamlines alignment-based searches and motif mining across large genomic datasets. SeqForge automates BLAST+ database creation and querying, integrates amino acid motif discovery, enables sequence and contig extraction, and curates results into structured, easily parsed formats. The platform supports diverse input formats, parallelized execution for high-performance computing environments, and built-in visualization tools. Benchmarking demonstrates that SeqForge achieves near-linear runtime scaling for computationally intensive modules while maintaining modest memory usage.

**Conclusions:** SeqForge lowers the computational barrier for large-scale meta/genomic exploration, enabling researchers to perform population-scale BLAST searches, motif detection, and sequence curation without custom scripting. The toolkit is freely available and platform-independent, making it suitable for both personal workstations and high-performance computing environments.

## Background

The rapid expansion of high-throughput sequencing has ushered in a new era of genomics, with thousands of microbial genomes and metagenomic assemblies deposited into public repositories each year. This expansion enables large-scale comparative analyses, genome mining initiatives, and meta-studies that uncover novel biological insight. Population-scale searches—such as pangenome exploration, biosynthetic gene cluster discovery, and antimicrobial resistance gene surveys—are increasingly essential for linking genomic diversity to ecological or clinical outcomes, as well as to drug discovery and development. However, leveraging this wealth of data requires computational tools that are both powerful and accessible to researchers with diverse technical expertise.

Numerous bioinformatics platforms support specific aspects of genome analysis. Quality assessment tools (e.g., QUAST [1], SeqKit2 [2], CheckM [3]), phylogenomic pipelines (e.g., PhyloPhlAn [4], getphylo [5], autoMLST [6], RAxML [7]), annotation frameworks (e.g., Anvi’o [8], eggNOG-mapper [9], AMRFinderPlus [10], Prokka [11]) and biosynthetic gene cluster discovery platforms (e.g., antiSMASH [12], ARTs [13], PRISM [14], RODEO [15]) allow researchers to assemble, annotate, and interpret genomic datasets. However, none of these platforms directly address a core need in exploratory meta/genomics: scalable, customizable alignment-based searches across large genome collections.

NCBI BLAST+ [16] remains the most widely used tool for sequence alignment and exploratory genome interrogation due to its robustness, speed, and versatility. However, population-scale studies with BLAST+ typically require custom scripting in Bash, Python, or similar languages to automate iterative database creation and query execution. For many researchers, particularly those without computational training, this requirement poses a significant barrier. Additionally, standard BLAST+ workflows generate a separate output file for each query–database comparison, making downstream organization, parsing, and interpretation cumbersome, even for moderately sized datasets.

To address these challenges, we developed SeqForge, a scalable command-line toolkit that streamlines large-scale genomic exploration. SeqForge automates key components of the BLAST+ workflow, simplifies the management of population-scale datasets, and adds features for amino acid motif discovery, sequence extraction, and organized results curation. By bridging the gap between existing pipelines and user-friendly exploratory search tools, SeqForge lowers the computational barrier for meta/genomic research and accelerates the interrogation of large genomic collections.

## Implementation

SeqForge is implemented in Python (>=3.10) and structured as a modular command-line toolkit with a centralized control unit. The package contains two core modules (Genome Search and Sequence Investigation) and a Utilities module (Figure 1). Each functional unit (e.g., makedb, query, extract) is implemented as a subcommand executed via a unified entry point (seqforge <subcommand> <args>), enabling access to all functions through a single interface. To maintain stability, each module is dynamically imported at runtime, so that failure in one component does not affect the operability of others. A built-in --module-health diagnostic command allows users to assess the availability of individual modules prior to execution. Internally, each module is structured around clear separation of responsibilities: input validation, multiprocessing logic, and output formatting. This modular design allows users to integrate individual components into custom workflows or use the complete SeqForge suite for full-scale data mining and exploration.

**Figure 1.**
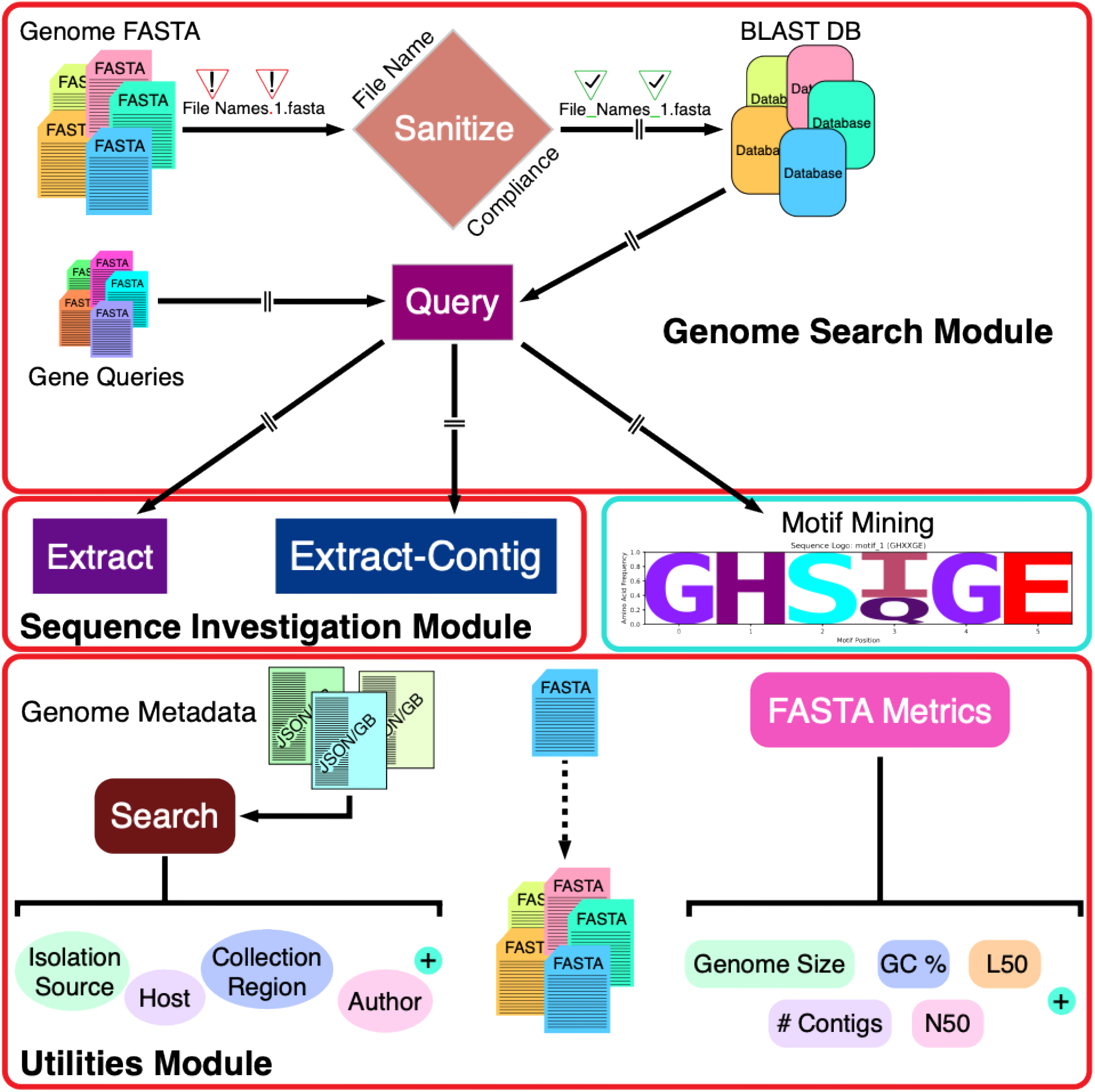
Overview of the SeqForge platform. The platform is organized into three modules: Genome Search Module includes database creation (makedb), database query (query), and motif mining; Sequence Investigation module includes extraction of aligned sequences (extract) and extraction of full contigs (extract-contig); Utilities Module includes metadata mining (search), FASTA file splitting (split-fasta), and calculation of assembly metrics (fasta-metrics). All modules are accessible via a unified command-line interface.

A unified file-handling system supports a wide range of FASTA input formats across all modules. This system allows for the submission of 1) individual FASTA files (including gzip-compressed formats), 2) directories containing multiple FASTA files, or 3) compressed archives (.zip, .tar, .tar.gz, .tgz) containing multiple FASTA files. Upon execution, SeqForge recursively enumerates and validates all FASTA files. For compressed archives, contents are extracted to a temporary working directory, which is automatically removed at the end of analysis unless the user specifies --keep-temp-files.

SeqForge was designed to streamline analysis of large meta/genomic datasets. For scalable performance, it uses Python’s concurrent.futures module for parallel execution. All computationally intensive modules (makedb, query, and extract) support multi-core processing via a --threads flag, allowing users to fully leverage available computational resources on personal laptops or high-performance computing (HPC) systems. SeqForge depends on BLAST+ 2.16.0, Biopython 1.85, pandas 2.3.1, matplotlib 3.10.5, seaborn 0.13.2, logomaker 0.8.7 [17], and numpy 2.3.2. All Python dependencies can be installed via pip, while BLAST+ is available through Conda or the NCBI FTP site. SeqForge is platform-independent and has been tested on Linux Ubuntu (v24.04) and macOS (v12.7.6) systems. All modules operate via the command line and are compatible with both personal workstations and HPC environments.

The core SeqForge workflow begins with creating a library of BLAST databases. Input file names are first scanned for special characters (e.g., non-extension periods, spaces, parentheses, etc.) that are incompatible with the BLAST+ architecture. If non-compliant characters are detected, SeqForge prints an error message and exits the program. Users can sanitize file names in two ways (excluding compressed archive submission): use the -- sanitize flag within the makedb command string to remove special characters in-place and proceed with database creation, or run the ‘seqforge sanitize’ utility script, specifying either in-place sanitization or copying files to a new directory and renaming them with the --sanitize-outdir <directory> option. Database type (nucl or prot) is determined automatically during database creation. Canonical nucleotide FASTA extensions (.fa, .fas, .ffn, .fna, .fasta) are parsed as nucleotide, while amino acid FASTA (.faa) are parsed as amino acid.

Next, users may proceed to the query module, which allows users to run any number of queries against any number of databases. First, database type is detected by the query module and execution of translated nucleotide (tblastn) or protein (blastp) query is automatically performed (nucleotide database + amino acid query = tblastn, protein database + amino acid query = blastp). Users can run nucleotide-nucleotide (blastn) searches by adding the --nucleotide-query flag. BLAST+ integration is achieved through subprocess calls to blastn, blastp, and tblastn, ensuring full compatibility with NCBI’s established alignment tools. SeqForge does not currently bundle BLAST+ binaries but is compatible with standard BLAST+ installations from Bioconda or from the NCBI FTP site.

By default, the query pipeline uses internally curated thresholds of 90% identity, 75% query coverage, and an E-value of 1 × 10^−5^. These values can be modified via command-line arguments. The standard output includes two concatenated results tables: all_results.csv, which contain all BLAST hits, regardless of inclusion criteria, and all_filtered_results.csv, which contain only hits meeting the percent identity, query coverage, and E-value thresholds. These tables include all standard BLAST+ fields, including qseqid, sseqid, pident, evalue, length, mismatch, gapopen, qstart, qend, sstart, send, bitscore, qlen, and sframe, along with calculated query coverage. SeqForge also generates standard BLAST+ alignment files for each query unless the user specifies --no-alignment-files. To return the strongest match per query per genome, the --report-strongest-matches flag can be employed, producing filtered_results.csv. Parallelization is implemented by assigning each query file as a separate task (i.e., one query file = one task). For optimal performance, each query file should harbor only one sequence. If a query file contains multiple sequences, the ‘seqforge split-fasta’ utility can split it into individual files. Multi-FASTA files can be used directly in the query pipeline; however, they are processed as a single task, which increases runtime.

The query module includes a motif mining function that parses blastp results for specific or loosely defined amino acid motifs via the --motif argument. Users may submit a single motif or space-separated list of motifs using the standard single-letter amino acid abbreviation, with ‘X’ serving as a wild card (e.g., WXWXIP). This option executes a modified regex search across all BLAST hits, regardless of inclusion thresholds, enabling detection within heterogenous gene families. Motifs can be linked to specific query files by appending the query base name (file name without extension) in braces within the command string (e.g., for ‘query_file.faa’: --motif <motif>{query_file}), which restricts parsing to results from that file. To facilitate downstream analysis, motif matches can be exported in FASTA format as either the full gene/domain alignment (--motif-fasta-out), or the specific motif string (--motif-fasta-out --motif-only). When > 1 motif is specified, SeqForge generates one FASTA file per motif.

Query results can be visualized by adding the --visualize flag to the query command string, with the standard output being a PNG at 300 dpi (output to PDF via --visualize --pdf). For queries against > 1 database, a heatmap will be generated (Figure 2A), with the color of each cell reflecting individual percent identity values per query per genome. For queries including motif searches, a sequence logo will additionally be generated using Logomaker [17] if any hits were returned (Figure 2B). Similar to the FASTA out option, for motif queries of > 1 unique motif string, a sequence logo will be generated for each search string.

**Figure 2.**
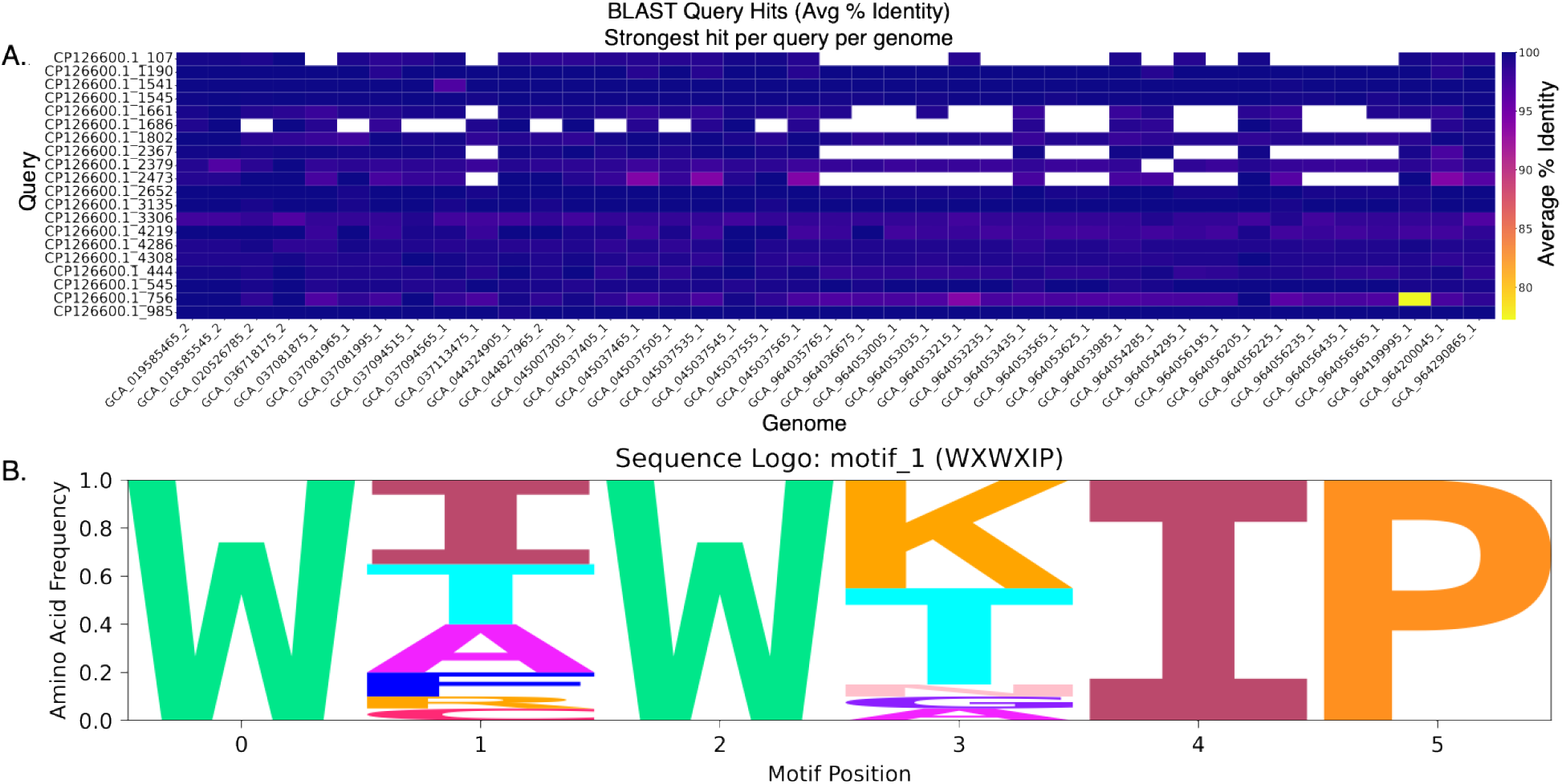
Example visualization output of the SeqForge query pipeline. **A** Heatmap generated using the --visualize option with 20 protein queries against 50 *E. coli* nucleotide databases. **B** Sequence logo generated using the -- motif WXWXIP –visualize command. Sequence logos were created with Logomaker [17]. Figures were minimally edited in Adobe Illustrator for clarity.

For downstream sequence analysis and annotation, SeqForge provides two extraction options for extraction of sequences identified via the query module: ‘seqforge extract’ and ‘seqforge extract-contig’. Users may pass a query results file (all_results.csv, all_filtered_results.csv, or filtered_results.csv) to the extract module, as well as the FASTA directory used to generate the BLAST database(s). Using ‘seqforge extract’, aligned sequences will be extracted to FASTA (multi-FASTA for > 1 hit) based on the alignment start and end values. For ‘seqforge extract’, up and/or downstream base pair padding may be specified to extract the aligned sequence plus *n* neighboring base pairs for more in-depth investigation of genomic context. Furthermore, as metagenome assemblies are often too large to open in their entirety using a standard genome browser, SeqForge additionally offers the extract-contig option, whereby entire contigs may be extracted based on a BLAST hit, facilitating manual sequence investigation using a genome browser.

SeqForge also includes lightweight utility modules for metadata compilation and assembly statistics. The first of these is ‘seqforge search’, which extracts metadata from NCBI-generated GenBank and/or JSON files, including accession number, organism, host, isolation source, and collection region. Users can extract all available metadata (--all) or select specific fields (--fields <fields>). Output formats include CSV, TSV and JSON. Missing metadata are reported as ‘not_specified’. This module supports both single-file or batch submission. Second, SeqForge offers the fasta-metrics tool, which quickly calculates common assembly stats, including genome size, number of contigs, contig length distribution, longest/shortest contig, GC %, N count, N50/N90, L50/L90, and the lengths of each contig as a ‘|’ separated list.

Example *Escherichia coli* (Table S1), *Streptomyces* (Table S2), and *Penicillium* (Table S3) genomes used for benchmarking were downloaded using NCBI Datasets [18]. Only complete genomes were included using the -- assembly-level complete Datasets argument. For *E. coli*, only assemblies released after March 1, 2024, were selected. Query sequences for the *E. coli* cohorts were generated by predicting coding sequences (CDS) from *E. coli* strain Ec1119 (Accession: GCA_008364465.2) with prodigal [19], producing a multi-FASTA amino acid file. This file was subsequently split into individual FASTA files using the SeqForge split-FASTA utility function, resulting in ∼4,400 predicted CDSs. From this population, cohorts of query sequences were randomly selected using an in-house Python script (20 or 50) to serve as query files. The tryptophan halogenase, RebH (Accession: Q8KHZ8), was downloaded from UniProt and used as the query to generate Figure 2. The *Penicillium*-trained CDS prediction model was generated using BRAKER2 [20], BUSCO (https://github.come/metashot/busco), RepeatModeler [21], and AUGUSTUS [22] (see Extended Methods for training protocol).

## Results

### Benchmarking and Performance

SeqForge is designed to expedite genome mining across datasets of any size, supporting batch submissions while maintaining flexibility and ease of interpreting large results datasets. To the best of our knowledge, there are no publicly available tools directly comparable to SeqForge. While some similar utility functions exist in QUAST [23] and SeqKit2 [2], these platforms focus on assembly and read quality analysis and sequence conversion and manipulation, rather than large-scale genome mining and analysis.

Benchmarking was performed for the makedb, query, extract/extract-contig modules to assess runtime scaling, memory requirements, and parallelization efficiency. Tests were performed on two representative datasets: 1) a “laptop-scale” dataset comprising 500 *E. coli* genomes and 20 gene queries, and 2) a “population-scale” dataset of 2,157 *E. coli* genomes with 50 gene queries, representing a typical workload for HPC environments. Tests used varying thread counts: 1, 8, and 16 threads for the 500-genome dataset; and 8, 16, 32, and 48 threads for the 2,157-genome dataset. Resource utilization was measured with ‘/usr/bin/time -v’, which reports wall clock runtime, average CPU usage, and maximum resident set size (RSS). Maximum RSS here reflects the peak memory used by any individual worker process, not the sum across all threads.

Across all modules, Seqforge demonstrated efficient scaling with increasing thread counts, with near-linear reductions in wall clock runtime for computationally intensive steps (makedb, query, and extract) (Table 1). For example, query execution on 500 genomes with 20 queries decreased from 32 minutes 32 seconds (1 thread) to 2 minutes using 16 threads. With the 2,157-genome, 50-query dataset, runtime dropped from 57 minutes 57 seconds (8 threads) to 15 minutes 15 seconds using 48 threads. Memory usage was modest for makedb (≤ 82 MB peak RSS) and moderate for query (≤ 901 MB peak RSS). Sequence extraction showed slightly diminishing returns beyond 16 threads, likely due to partial I/O-bound performance, but remained efficient, with extraction of > 100,000 sequences completing just over 3 minutes using 8 threads and just over 1 minute at 48 threads (Table 1).

**Table 1.** Multiprocessing performance and resource usage of SeqForge modules. Wall clock runtimes and peak memory usage (maximum resident size, RSS, in megabytes) for makedb, query, extract, and extract-contig modules. These times are reported as minutes:seconds (m:s). RSS reflects the maximum memory used by any single worker process, not the sum across threads.

| Module | # Genomes | # Queries | # Sequences | # Threads | Runtime (m:s) | RSS (MB) | Avg. CPU (%) |
| --- | --- | --- | --- | --- | --- | --- | --- |
| Makedb | 500 | N/A | N/A | 1 | 0:47 | 79 | 107 |
|  |  |  |  | 8 | 0:06 | 79 | 808 |
|  |  |  |  | 16 | 0:03 | 80 | 1524 |
|  | 2157 | N/A | N/A | 8 | 0:23 | 81 | 786 |
|  |  |  |  | 16 | 0:12 | 81 | 1537 |
|  |  |  |  | 32 | 0:07 | 81 | 2974 |
|  |  |  |  | 48 | 0:05 | 82 | 4411 |
| Query | 500 | 20 | N/A | 1 | 32:33 | 115 | 100 |
|  |  |  |  | 8 | 3:55 | 114 | 790 |
|  |  |  |  | 16 | 2:00 | 113 | 1558 |
|  | 2157 | 50 | N/A | 8 | 57:57 | 901 | 667 |
|  |  |  |  | 16 | 22:07 | 885 | 898 |
|  |  |  |  | 32 | 16:37 | 885 | 1253 |
|  |  |  |  | 48 | 15:15 | 885 | 1619 |
| Query + Motif | 150 | 1 | N/A | 1 | 0:41 | 1565 | 110 |
|  |  |  |  | 8 | 0:18 | 1565 | 315 |
|  |  |  |  | 16 | 0:34 | 1569 | 261 |
|  |  | 2 | N/A | 1 | 5:09 | 1597 | 101 |
|  |  |  |  | 8 | 1:07 | 1578 | 543 |
|  |  |  |  | 16 | 0:48 | 1578 | 793 |
|  |  | 4 | N/A | 1 | 14:38 | 1661 | 100 |
|  |  |  |  | 8 | 2:28 | 1601 | 698 |
|  |  |  |  | 16 | 1:33 | 1597 | 1136 |
| Extract | 500 | 20 | 10,003 | 1 | 2:27 | 1225 | 104 |
|  |  |  |  | 8 | 0:21 | 1117 | 815 |
|  |  |  |  | 16 | 0:11 | 1063 | 1583 |
|  | 2157 | 50 | 100,524 | 8 | 3:09 | 1138 | 808 |
|  |  |  |  | 16 | 1:49 | 1134 | 1550 |
|  |  |  |  | 32 | 1:15 | 1129 | 2674 |
|  |  |  |  | 48 | 1:14 | 1094 | 2908 |
| Extract-Contig | 500 | 20 | 500 | 1 | 0:17 | 9947 | 124 |
|  |  |  |  | 8 | 0:18 | 10156 | 126 |
|  |  |  |  | 16 | 0:18 | 9769 | 127 |
|  | 2157 | 50 | 2145 | 8 | 0:59 | 33734 | 115 |
|  |  |  |  | 16 | 1:02 | 27705 | 114 |
|  |  |  |  | 32 | 0:56 | 14448 | 116 |
|  |  |  |  | 48 | 0:57 | 14773 | 117 |

In contrast, extract-contig showed minimal speedup with more threads, consistent with an I/O-bound workload dominated by sequential file reads and writes. Peak RSS reached up to 33 GB when processing the 2,157-genome dataset, due to loading and writing entire contigs, though runtime remained under 1 minute even at large scale. Because most genomes in the benchmark dataset were complete assemblies with < 8 contigs, many contig extractions involved extracting whole chromosomal sequences. As a result, contig extraction was likely dominated by disk I/O rather than CPU processing and exhibited little benefit from parallelization compared to sequence extraction, which operates on smaller regions. Overall, SeqForge maintains modest per-thread memory usage, scales efficiently for computationally intensive modules, and delivers fast performance even for I/O-heavy workflows.

### Motif Mining

To demonstrate Seqforge’s motif mining capabilities, we conducted a targeted search for canonical catalytic and stereochemistry-associated motifs within a well-characterized biosynthetic gene cluster (BGC). Specifically, we mined the erythromycin BGC (MIBiG accession: BGC0000055) for domain-level motifs associated with acyltransferase (AT), ketosynthase (KS), and ketoreductase (KR) activity across the polyketide synthase (PKS) modules. The erythromycin BGC consists of a loading module plus six chain-elongation modules containing seven AT domains (substrate specificity), six KS domains (condensation activity), six KR domains (reduction and stereochemistry determination), one dehydratase (DH) and one enoyl reductase (ER) domain [24, 25]. While the ER domain plays a role in final stereochemistry of the C-2 position—particularly the Y/V52’ residue [25, 26]— this benchmark focused solely on AT, KS, and KR motifs responsible for upstream chain elongation and modification of the C-2 and β-keto positions of the polyketide backbone. Motif search strings were defined using common PKS domain motifs for malonyl and methylmalonyl substrate selectivity, KS catalytic activity, and KR- associated stereochemistry determination [25, 27, 28]. The motif analysis was executed using the following command string: ‘--motif RVXXXQ{AT} GHXXGE{AT} YXXH{AT} HXSH{AT} HXFH{AT} TAXSSX{KS} HXAXXLDDX{KR} SSXXXXXXXXXXXXYXX{KR}’. SeqForge returned results as a structured table for all PKS modules (Table S4) and successfully detected the expected AT, KS, and KR motifs across all relevant modules (Table 2, Figures S1–S3). Importantly, motif parsing occurs within the context of full alignment sequence, enabling researchers to extract matches directly using the --motif-fasta-out argument for downstream analysis. These results align with previously characterized enzymatic roles for each biosynthetic module, validating the utility of motif mining as a complementary tool for BGC functional annotation.

**Table 2.**
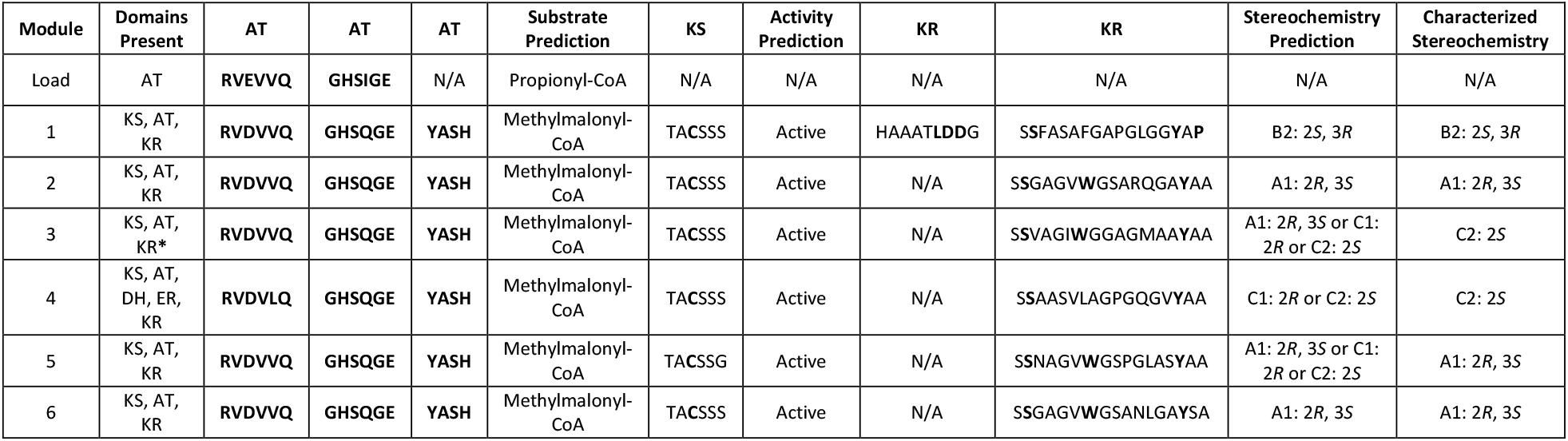
Domain presence, substrate specificity, activity prediction and extracted amino acid motifs from the erythromycin biosynthetic gene cluster (BGC0000055). Domain composition, predicted substrate specificity, and detected motifs for acyltransferase (AT), ketosynthase (KS), and ketoreductase (KR) domains identified using SeqForge’s --motif function with domain-linked search strings. KR* indicates an inactive ketoreductase domain [25]. N/A: no motif hits detected. Key residues are in bold.

### Leveraging SeqForge to Detect Potential Copper-Dependent Halogenases Across *Penicillium*

To demonstrate the broader utility of SeqForge, we applied the full pipeline to an uncharacterized fungal population. As a case study, we selected ApnU, a recently characterized copper-dependent halogenase from the *Penicillium oxalicum* 114-2 atpenin B BGC (MIBiG accession: BGC0002067) (Table S5) [29]. Unlike Fe(II)/α- ketoglutarate-dependent halogenases, ApnU catalyzes mono- or iterative chlorination of unactivated C(*sp*^*3*^)–H bonds via a unique pair of HXXHC motifs responsible for coordination of Cu(II), and it can also incorporate other halides or pseudohalides [29]. Given its mechanistic novelty, ApnU presented a compelling target for mining functionally analogous or homologous BGCs across the *Penicillium* genus. A dataset of 549 publicly available *Penicillium* genomes was curated from NCBI Datasets (Table S3) and converted to BLAST databases using ‘seqforge makedb’. A translated nucleotide search was performed with ‘seqforge query’, using the ApnU amino acid sequence as input and a relaxed inclusion threshold of ≥ 80% identity and ≥ 70% query coverage. Fourteen hits met these criteria and were retained for downstream analysis (Table S6).

Genomic context was assessed by extracting each hit along with 50 kb upstream and 30 kb downstream flanking sequences using ‘seqforge extract’. Coding sequences were predicted with AUGUSTUS, trained on the high-quality *P. chrysogenum* IBT 35668 genome (GCA_028827035.1), and validated against the canonical atpenin B cluster. Protein databases from these predictions were analyzed using SeqForge’s motif mining utility, which identified exactly two HXXHC motifs in each case, consistent with the ApnU copper-binding residues. To confirm the accuracy of motif detection, hits were extracted and aligned with the canonical ApnU sequence using MUSCLE [30]. Both HXXHC motifs were confirmed in all hits (Figure 3A, Figure S4), supporting the conservation of the copper-binding residues.

**Figure 3.**
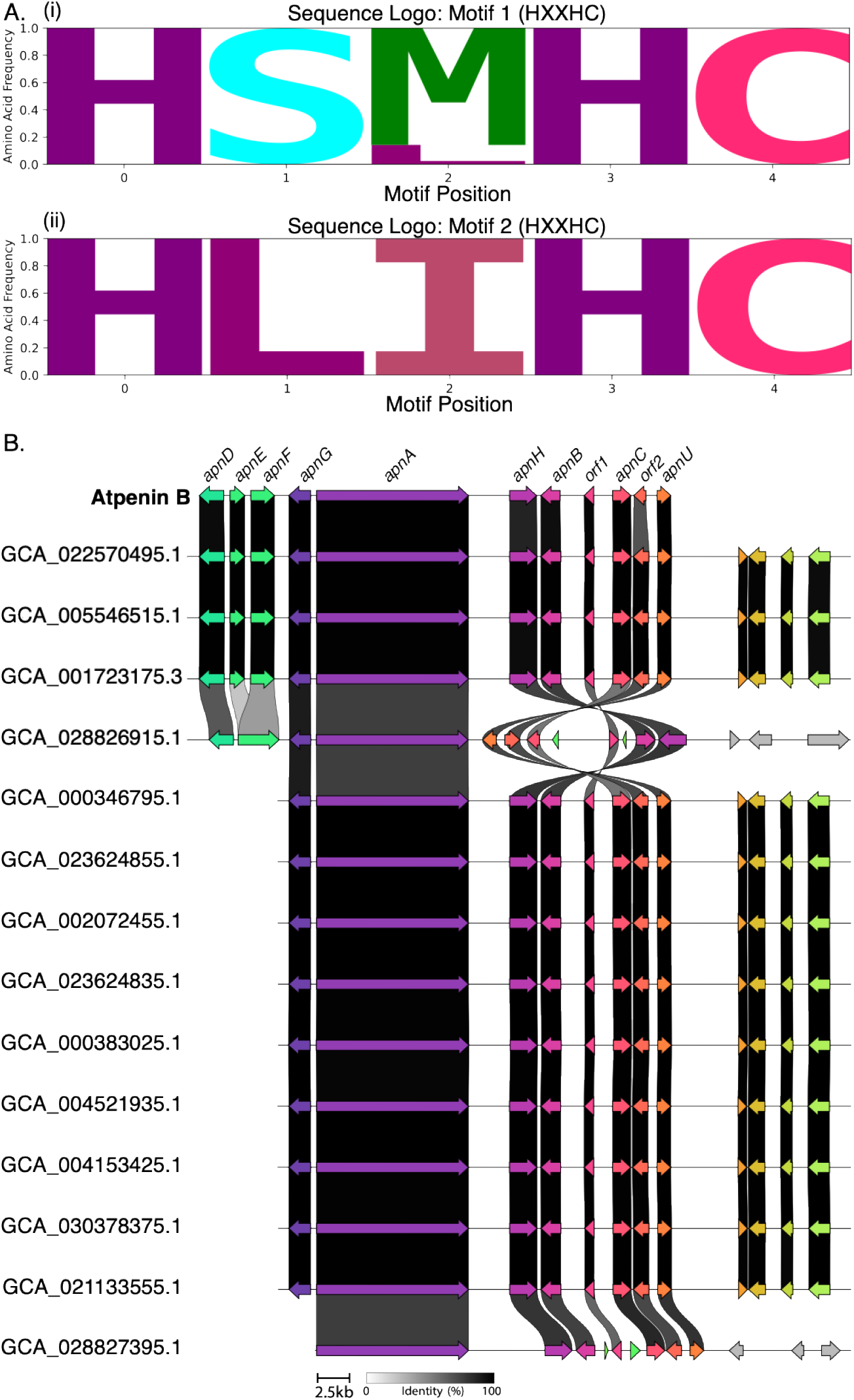
Atpenin B-like biosynthetic gene clusters (BGCs) identified in publicly available *Penicillium* genomes using the full SeqForge analytical pipeline. **A** The first (i) and second (ii) HXXHC copper-binding motifs required for catalytic activity of the ApnU copper-dependent halogenase. Sequence logos represent predicted ApnU homologs. **B** Synteny map of atpenin B-like BGCs identified in publicly available *Penicillium* genomes. orf1 and orf2 represent genes within the atpenin B BGC that have not yet been formally characterized [32].

To confirm that these genomic regions encoded atpenin B-like clusters, each sequence and its AUGUSTUS-generated GFF3 file were submitted to antiSMASH’s fungal pipeline [12]. All 14 regions showed medium-to-high similarity to the characterized atpenin B gene cluster (Figure 3B). Clinker alignments via CAGECAT [31] revealed strong synteny and high gene identity across all hits. Collectively, this analysis demonstrates SeqForge’s end-to-end capability for population-scale homology screening, flanking region extraction, motif detection, and downstream BGC characterization, enabling rapid discovery of biosynthetic diversity from large genome datasets.

### Utilities

Seqforge includes several auxiliary utilities designed to streamline genome mining workflows. The ‘split-fasta’ command fragments large multi-FASTA files into either single-record files or user-defined batches of *n* sequences. The ‘search’ module extracts genomic metadata (e.g., accession, isolation source, host, and related descriptors) from GenBank and JSON files and was used here to compile metadata for the test populations (Tables S1–S3). For rapid quality checks, the lightweight ‘fasta-metrics’ module reports basic assembly statistics and was benchmarked against QUAST on three *E. coli* and three *Streptomyces* genomes (Table S7).

Many annotation and CDS-prediction pipelines assign generic identifiers (e.g., “hypothetical protein”) or placeholder headers (e.g., from AUGUSTUS), which can result in duplicate sequence IDs within or across files. Such duplications can disrupt downstream parsing and interfere with SeqForge’s motif mining workflow. To address this, SeqForge provides a ‘unique-header’ utility that ensures collision-free FASTA identifiers across inputs. This tool preserves the original label while appending the source filename and a short alphanumeric tag.

For example: >hypothetical (from genome1a.faa) becomes >hypothetical_genome1a_54uMe. This approach maintains identifiability across large, multi-file datasets and reduces the risk of bookkeeping or programmatic errors during downstream analyses.

## Discussion

SeqForge addresses a practical gap between small BLAST+ searches and population-scale mining by providing a modular command-line interface that automates database construction, high-throughput querying, motif discovery, and downstream extraction/visualization. In contrast to pan-genome pipelines, HMM-centric profilers, or general QC/formatting toolkits, SeqForge preserves the interpretability and ubiquity of NCBI BLAST+ while adding scalability, uniform outputs, and utilities that smooth common stress points (e.g., filename sanitization, header de-duplication, and lightweight assembly metrics). This design lowers the barrier for exploratory meta/genomic analyses.

While SeqForge addresses a clear gap in currently available software, it does have limitations. Results remain sensitive to BLAST thresholds and database quality. Permissive settings can inflate false positives for promiscuous families, whereas overly stringent cutoffs may exclude divergent homologs. Dynamic querying, such as using multiple representative sequences or varying inclusion thresholds across runs, can help balance these trade-offs and strengthen result datasets. Similarly, the regex-based motif search, while fast, transparent, and well-suited for highly conserved motifs, may overlook gapped or degenerate variants that could be more effectively captured using profiles or HMMs.

## Conclusion

The field of microbial genomics has made great strides in developing computational platforms that enrich microbial discovery, characterization, annotation, and functional investigation. SeqForge builds on this progress by providing an accessible, scalable solution for performing multi-query BLAST searches and motif mining across large genomic datasets. By integrating these capabilities into a streamlined workflow, SeqForge reduces the need for custom scripting, minimizes manual curation, and accelerates the identification of conserved functional motifs. Its modular design and support for multi-core execution make it equally suitable for small-scale analyses on personal computers and high-throughput screening on HPC clusters. SeqForge is freely available and adaptable to a wide range of genomic mining applications, helping accelerate the pace of discovery from the ever-growing wealth of publicly available genome datasets.

## Availability and Requirements

**Project name:** SeqForge

**Home page:** at https://github.com/ERBringHorvath/SeqForge

**Operating system:** Platform independent

**Programming language:** Python

**License:** MIT

**Any restrictions to use by non-academics** None.

## Abbreviations

AT: acyltransferase
BGC: biosynthetic gene cluster
CDS: Coding sequences
DH: dehydratase
ER: enoylreductase
HMM: hidden Markov model
HPC: High performance computing
KR: ketoreductase
KS: ketosynthase
PKS: polyketide synthase
RSS: resident set size

## Declarations

### Ethics approval and consent to participate

Not applicable

## Consent for publication

Not applicable.

## Availability of data and materials

SeqForge is available through GitHub at https://github.com/ERBringHorvath/SeqForge. Please read the SeqForge documentation for information on installation and usage.

## Authors’ contributions

E.R.B.H. wrote code, designed and performed experiments, and conceived the project.

J.M.W. helped design experiments and supervised the project. All authors wrote and reviewed the manuscript.

## Acknowledgements

We thank Mathew Stein for valuables discussion on the design of SeqForge and for reviewing the draft manuscript.

