## Supplemental Information for "SeqForge: A scalable platform for alignment-based searches, motif detection, and sequence curation across meta/genomic datasets"

**Table of Contents**

Extended Methods 2
Tables S1-S3: Information on genomes used in this study. 3
Table S4: Erythromycin motif mining output 4
Figure S1: Aligned erythromycin AT domains 6
Figure S2: Aligned erythromycin KS domains 7
Figure S3: Aligned erythromycin KR domains 8
Table S5: Predicted functions of atpenin B biosynthetic genes 9
Table S6: *Penicillium* genomes meeting ApnU tblastn inclusion thresholds 10
Figure S4: Aligned ApnU sequences 11
Table S7: SeqForge FASTA-metrics module compared to QUAST assembly metrics output 12
References 14

### Extended Methods

Augustus Penicillium Model

The genome and predicted coding sequences of the *Penicillium chrysogenum* strain IBT 35668 (Accession: GCA_028827035.1) was downloaded from NCBI and used as a reference for model training. Training and coding sequence (CDS) prediction was carried out using Braker2 v2.1.6 [1], BUSCO v5.4.3 (https://github.com/metashot/busco), RepeatModeler v1.0.8 [2], and AUGUSTUS v3.5.0 [3]. RepeatModeler was run using default settings and returned an empty consensus file; upon further investigation, the *P. chrysogenum* IBT 35668 genome had already undergone soft masking. BRAKER2 was executed using the following command string: braker.pl --genome ./genome/GCA_028827035_1.fna --prot_seq ./proteins/GCA_028827035_1_proteins.faa --softmasking --gff3 --cores 32. BUSCO was run as follows: busco -i braker/augustus.hints.aa -l eurotiomycetes_odb10 -m proteins -o busco_penicillium -c 16, and returned the following results: C:98.5% (S: 97.6%, D: 0.9%) F: 0.8%, M: 0.7%, n: 3546. Where: C = 3493 complete BUSCOS, S = 3462 complete and single-copy BUSCOS, D = 31 complete and duplicated BUSCOS, F = 27 fragmented BUSCOS, and M = 26 missing BUSCOS. CDS predictions were performed with AUGUSTUS using default settings.

Tables S1–S3: Information on the *E. coli*, *Streptomyces,* and *Penicillium* genomes used in this study. Due to their size, these tables are available as separate downloadable files.

Table S4. Raw output of key motif mining in erythromycin acyltransferase (AT), ketosynthase (KT), and ketoreductase (KR) domains. Motif 5 (HXFH) returned no hits, and those columns were therefore removed. A single off-target hit for HXSH was returned but discarded as it overlapped with the appropriate YXXH motif.

| genome | query | sseqid | sstart | send | motif_1 | motif_1_ pattern | motif_2 | motif_2_ pattern | motif_3 | motif_3_ pattern | motif_4 | motif_4_ pattern | motif_6 | motif_6_ pattern | motif_7 | motif_7_ pattern | motif_8 | motif_8_ pattern | match_ start | match _end |
|---|---|---|---|---|---|---|---|---|---|---|---|---|---|---|---|---|---|---|---|---|
| erythromycin _BGC | AT2 | erythromycin_S_ erythraea_13 | 2016 | 2311 | RVDVLQ | RVXXXQ | | | | | | | | | | | | | 2072 | 2077 |
| erythromycin _BGC | AT2 | erythromycin_S_ erythraea_11 | 1048 | 1348 | RVDVVQ | RVXXXQ | | | | | | | | | | | | | 1110 | 1115 |
| erythromycin _BGC | AT2 | erythromycin_S_ erythraea_11 | 2526 | 2822 | RVDVVQ | RVXXXQ | | | | | | | | | | | | | 2581 | 2586 |
| erythromycin _BGC | AT2 | erythromycin_S_ erythraea_13 | 562 | 832 | RVDVVQ | RVXXXQ | | | | | | | | | | | | | 618 | 623 |
| erythromycin _BGC | AT2 | erythromycin_S_ erythraea_14 | 556 | 830 | RVDVVQ | RVXXXQ | | | | | | | | | | | | | 609 | 614 |
| erythromycin _BGC | AT2 | erythromycin_S_ erythraea_14 | 2019 | 2309 | RVDVVQ | RVXXXQ | | | | | | | | | | | | | 2074 | 2079 |
| erythromycin _BGC | AT2 | erythromycin_S_ erythraea_11 | 73 | 366 | RVEVVQ | RVXXXQ | | | | | | | | | | | | | 126 | 131 |
| erythromycin _BGC | KR1 | erythromycin_S_ erythraea_11 | 1628 | 1805 | | | | | | | | | | | HAAATL DDG | HXAXXLD DX | | | 1713 | 1721 |
| erythromycin _BGC | KR1 | erythromycin_S_ erythraea_11 | 1628 | 1805 | | | | | | | | | | | | | SSFASAFG APGLGGY AP | SSXXXXXXX XXXXXYXX | 1761 | 1777 |
| erythromycin _BGC | KR1 | erythromycin_S_ erythraea_11 | 3073 | 3249 | | | | | | | | | | | | | SSGAGV WGSARQ GAYAA | SSXXXXXXX XXXXXYXX | 3205 | 3221 |
| erythromycin _BGC | KR1 | erythromycin_S_ erythraea_14 | 1116 | 1292 | | | | | | | | | | | | | SSNAGV WGSPGLA SYAA | SSXXXXXXX XXXXXYXX | 1248 | 1264 |
| erythromycin _BGC | KR1 | erythromycin_S_ erythraea_14 | 2556 | 2730 | | | | | | | | | | | | | SSGAGV WGSANL GAYSA | SSXXXXXXX XXXXXYXX | 2686 | 2702 |
| erythromycin _BGC | KR1 | erythromycin_S_ erythraea_13 | 3141 | 3317 | | | | | | | | | | | | | SSAASVLA GPGQGVY AA | SSXXXXXXX XXXXXYXX | 3273 | 3289 |
| erythromycin _BGC | KR1 | erythromycin_S_ erythraea_13 | 1132 | 1297 | | | | | | | | | | | | | SSVAGIW GGAGMA AYAA | SSXXXXXXX XXXXXYXX | 1253 | 1269 |
| erythromycin _BGC | AT2 | erythromycin_S_ erythraea_11 | 1048 | 1348 | | | GHSQGE | GHXXGE | | | | | | | | | | | 1141 | 1146 |
| erythromycin _BGC | AT2 | erythromycin_S_ erythraea_11 | 1048 | 1348 | | | | | YASH | YXXH | | | | | | | | | 1240 | 1243 |
| erythromycin _BGC | AT2 | erythromycin_S_ erythraea_11 | 1048 | 1348 | | | | | | | HSSH | HXSH | | | | | | | 1243 | 1246 |
| erythromycin _BGC | AT2 | erythromycin_S_ erythraea_11 | 2526 | 2822 | | | GHSQGE | GHXXGE | | | | | | | | | | | 2612 | 2617 |
| erythromycin _BGC | AT2 | erythromycin_S_ erythraea_11 | 2526 | 2822 | | | | | YASH | YXXH | | | | | | | | | 2716 | 2719 |
| erythromycin _BGC | AT2 | erythromycin_S_ erythraea_11 | 73 | 366 | | | GHSIGE | GHXXGE | | | | | | | | | | | 157 | 162 |
| erythromycin _BGC | AT2 | erythromycin_S_ erythraea_13 | 2016 | 2311 | | | GHSQGE | GHXXGE | | | | | | | | | | | 2103 | 2108 |

Table S4 continued.

| genome | query | sseqid | sstart | send | motif_1 | motif_1_ pattern | motif_2 | motif_2_ pattern | motif_3 | motif_3_ pattern | motif_4 | motif_4_ pattern | motif_6 | motif_6_ pattern | motif_7 | motif_7_ pattern | motif_8 | motif_8_ pattern | match_ start | match _end |
|---|---|---|---|---|---|---|---|---|---|---|---|---|---|---|---|---|---|---|---|---|
| erythromycin _BGC | AT2 | erythromycin_S_ erythraea_13 | 2016 | 2311 | | | | | YASH | YXXH | | | | | | | | | 2205 | 2208 |
| erythromycin _BGC | AT2 | erythromycin_S_ erythraea_13 | 562 | 832 | | | GHSQGE | GHXXGE | | | | | | | | | | | 649 | 654 |
| erythromycin _BGC | AT2 | erythromycin_S_ erythraea_13 | 562 | 832 | | | | | YASH | YXXH | | | | | | | | | 751 | 754 |
| erythromycin _BGC | AT2 | erythromycin_S_ erythraea_14 | 556 | 830 | | | GHSQGE | GHXXGE | | | | | | | | | | | 640 | 645 |
| erythromycin _BGC | AT2 | erythromycin_S_ erythraea_14 | 556 | 830 | | | | | YASH | YXXH | | | | | | | | | 742 | 745 |
| erythromycin _BGC | AT2 | erythromycin_S_ erythraea_14 | 2019 | 2309 | | | GHSQGE | GHXXGE | | | | | | | | | | | 2105 | 2110 |
| erythromycin _BGC | AT2 | erythromycin_S_ erythraea_14 | 2019 | 2309 | | | | | YASH | YXXH | | | | | | | | | 2207 | 2210 |
| erythromycin _BGC | KS | erythromycin_S_ erythraea_11 | 522 | 945 | | | | | | | | | TACSSS | TAXSSX | | | | | 690 | 695 |
| erythromycin _BGC | KS | erythromycin_S_ erythraea_11 | 1997 | 2416 | | | | | | | | | TACSSS | TAXSSX | | | | | 2162 | 2167 |
| erythromycin _BGC | KS | erythromycin_S_ erythraea_13 | 1491 | 1910 | | | | | | | | | TACSSS | TAXSSX | | | | | 1659 | 1664 |
| erythromycin _BGC | KS | erythromycin_S_ erythraea_13 | 33 | 457 | | | | | | | | | TACSSS | TAXSSX | | | | | 200 | 205 |
| erythromycin _BGC | KS | erythromycin_S_ erythraea_14 | 41 | 450 | | | | | | | | | TACSSG | TAXSSX | | | | | 197 | 202 |
| erythromycin _BGC | KS | erythromycin_S_ erythraea_14 | 1488 | 1909 | | | | | | | | | TACSSS | TAXSSX | | | | | 1655 | 1660 |

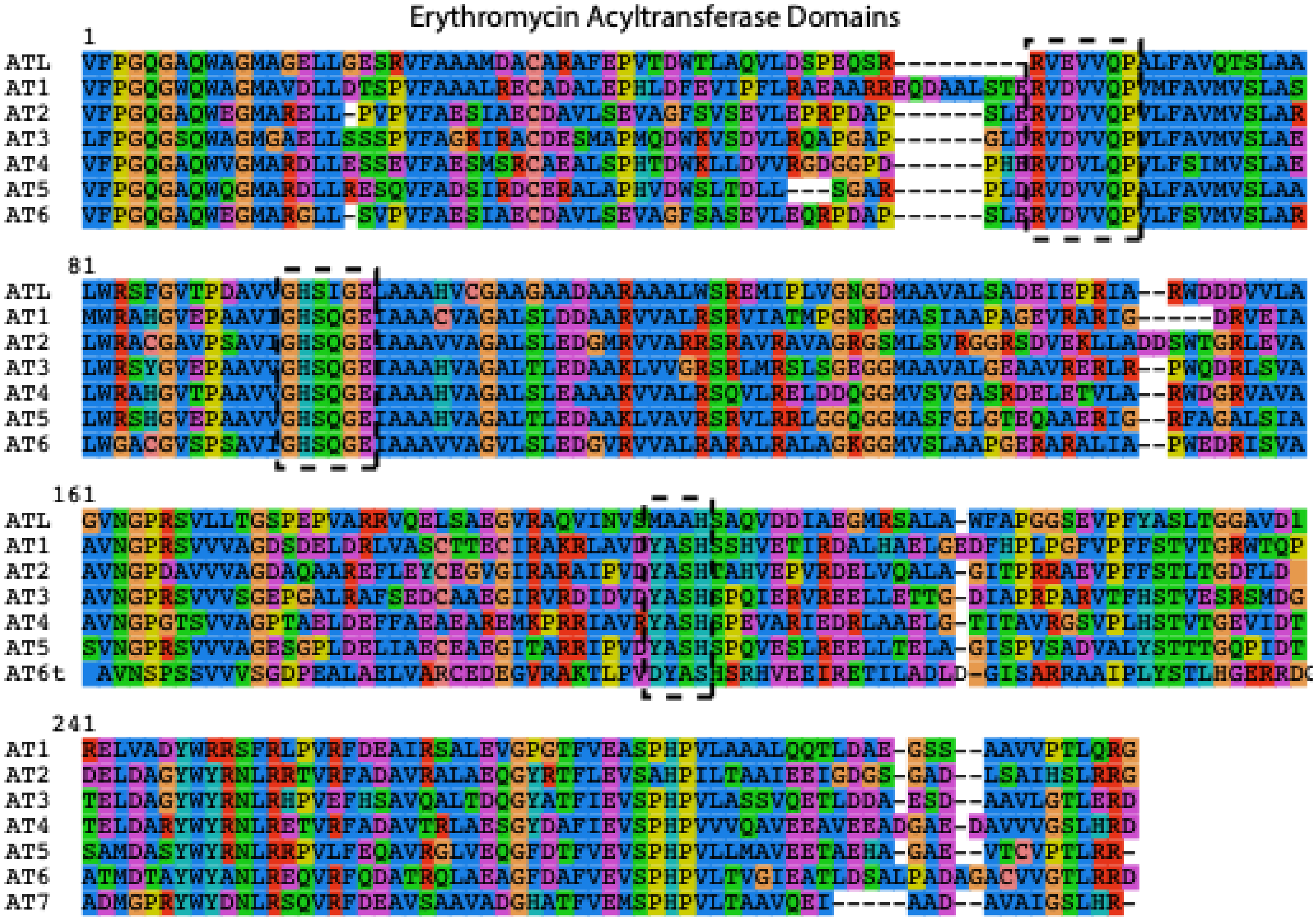

Figure S1. Aligned acyltransferase (AT) domains from the erythromycin biosynthetic gene cluster in *Saccharopolyspora erythraea* (MIBiG accession BGC0000055). ATL: acyltransferase loading domain. Key motifs are outlined with black dashed lines.

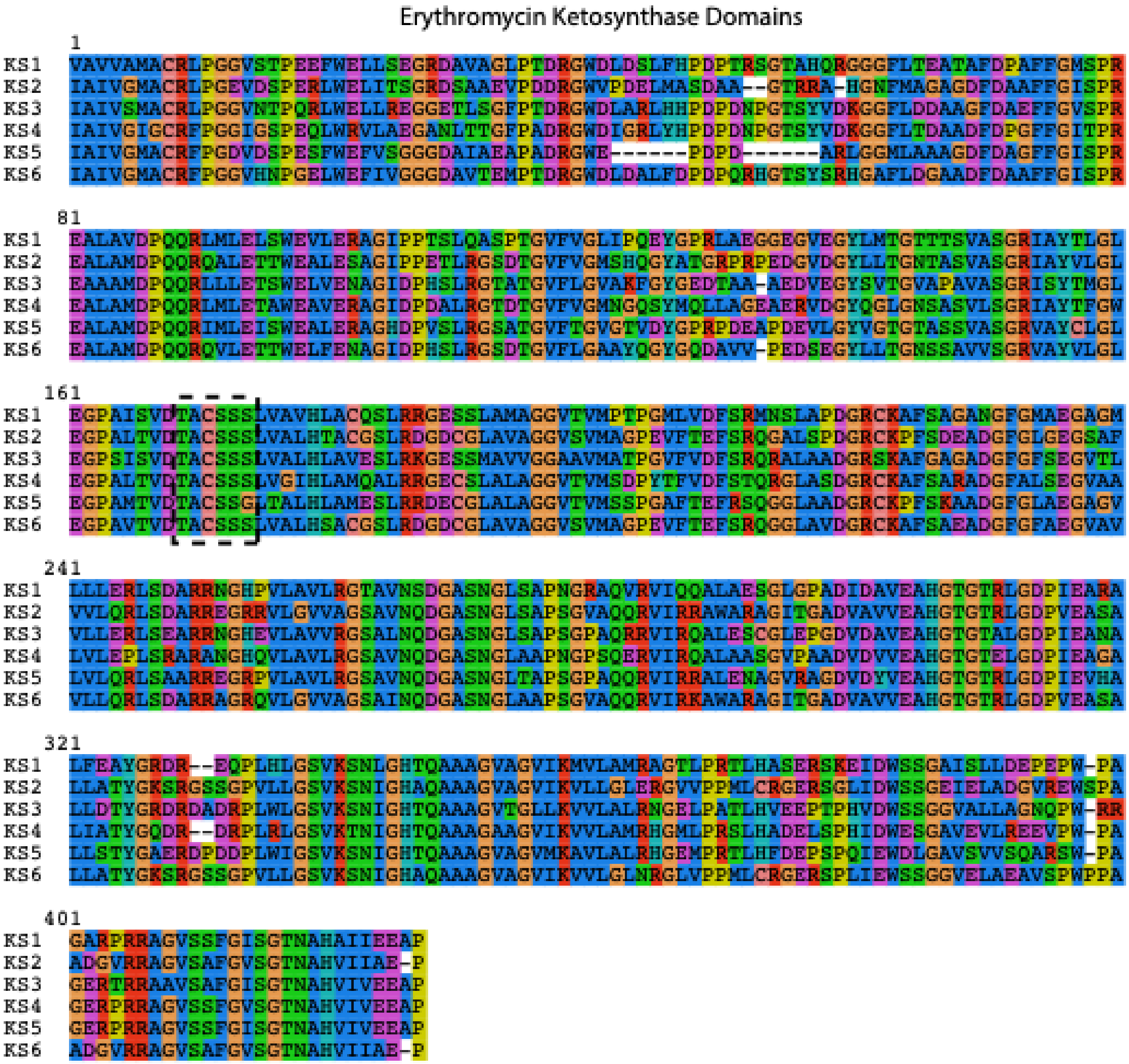

Figure S2. Aligned ketosynthase (KS) domains from the erythromycin biosynthetic gene cluster in *Saccharopolyspora erythraea* (MIBiG accession BGC0000055). Active site motif is outlined with a black dashed line.

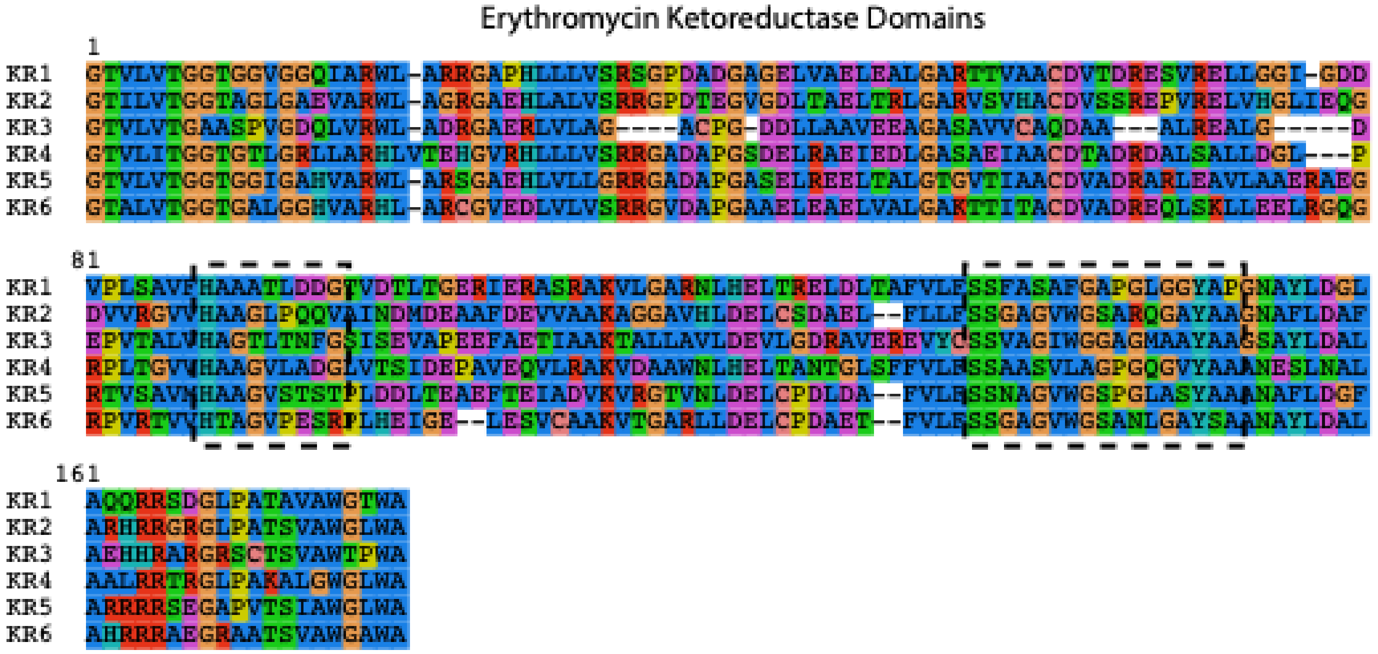

Figure S3. Aligned ketoreductase (KR) domains from the erythromycin biosynthetic gene cluster in *Saccharopolyspora erythraea* (MIBiG accession BGC0000055). Key motifs are outlined with black dashed lines.

Table S5. Predicted functions of atpenin B biosynthetic genes. *function has been biochemically characterized, but NCBI BLAST annotation has not been updated.

| Gene | Function/Predicted Function | Identity (%) | Query Coverage (%) | NCBI Accession |
|---|---|---|---|---|
| *apnU* | Cu(II)-dependent halogenase* | 100 | 100 | XP_049973506.1 |
| *orf2* | Hypothetical | 100 | 100 | EPS34229.1 |
| *apnC* | FAD-dependent monooxygenase | 100 | 99.3 | XP_049973504.1 |
| *orf1* | Hypothetical | 100 | 100 | EPS34231.1 |
| *apnB* | Hypothetical | 100 | 99.76 | XP49973502.1 |
| *apnH* | Hypothetical | 100 | 100 | EPS34233.1 |
| *apnA* | Polyketide synthase | 100 | 98.97 | XP049973499.1 |
| *apnG* | Cytochrome monooxygenase | 100 | 100 | XP049973498.1 |
| *apnF* | FAD-dependent monooxygenase | 100 | 99.83 | XP049973497.1 |
| *apnE* | Trans-enoyl reductase | 100 | 99.72 | XP049973496.1 |
| *apnD* | Cytochrome monooxygenase | 100 | 99.58 | XP_049973495.1 |

Table S6. *Penicillium* genomes meeting ApnU query inclusion thresholds of % identity ≥ 80% and query coverage ≥ 70%. Table represents all_filtered_results.csv generated from SeqForge's query module.

| qseqid | sseqid | database | query_file_name | pident | query_coverage | evalue | bitscore | length | mismatch | gapopen | qstart | qend | sstart | send | qlen | sframe |
|---|---|---|---|---|---|---|---|---|---|---|---|---|---|---|---|---|
| ApnU | JARG01000001.1 | GCA_002072455_1_Pexp1_0_genomic | ApnU | 84.775 | 99.59 | 9.69E-162 | 495 | 289 | 0 | 2 | 1 | 245 | 6402133 | 6401267 | 245 | -3 |
| ApnU | SDBR01000018.1 | GCA_004521935_1_ASM452193v1_genomic | ApnU | 84.775 | 99.59 | 9.25E-162 | 495 | 289 | 0 | 2 | 1 | 245 | 324004 | 323138 | 245 | -2 |
| ApnU | CP093053.1 | GCA_022570495_1_ASM2257049v1_genomic | ApnU | 84.775 | 99.59 | 9.51E-162 | 495 | 289 | 0 | 2 | 1 | 245 | 2315783 | 2316649 | 245 | 2 |
| ApnU | KB644415.1 | GCA_000346795_1_pde_v1_0_genomic | ApnU | 84.775 | 99.59 | 9.36E-162 | 495 | 289 | 0 | 2 | 1 | 245 | 3584465 | 3583599 | 245 | -2 |
| ApnU | JASKYK010000001.1 | GCA_030378375_1_ASM3037837v1_genomic | ApnU | 84.775 | 99.59 | 9.55E-162 | 495 | 289 | 0 | 2 | 1 | 245 | 3522649 | 3521783 | 245 | -2 |
| ApnU | CM041256.1 | GCA_001723175_3_ASM172317v3_genomic | ApnU | 84.775 | 99.59 | 9.55E-162 | 495 | 289 | 0 | 2 | 1 | 245 | 2334210 | 2335076 | 245 | 3 |
| ApnU | CP088335.1 | GCA_021133555_1_ASM2113355v1_genomic | ApnU | 84.775 | 99.59 | 9.47E-162 | 495 | 289 | 0 | 2 | 1 | 245 | 3546551 | 3545685 | 245 | -1 |
| ApnU | KB908904.1 | GCA_000383025_1_pdt_v1_0_genomic | ApnU | 84.775 | 99.59 | 9.52E-162 | 495 | 289 | 0 | 2 | 1 | 245 | 3534979 | 3534113 | 245 | -2 |
| ApnU | SZWD01000001.1 | GCA_005546515_1_SGAir0226_genomic | ApnU | 84.775 | 99.59 | 5.70E-162 | 496 | 289 | 0 | 2 | 1 | 245 | 1257011 | 1257877 | 245 | 2 |
| ApnU | JAMABO010000024.1 | GCA_023624835_1_ASM2362483v1_genomic | ApnU | 84.775 | 99.59 | 9.24E-162 | 495 | 289 | 0 | 2 | 1 | 245 | 195075 | 194209 | 245 | -1 |
| ApnU | JAQJAE010000006.1 | GCA_028827395_1_ASM2882739v1_genomic | ApnU | 81.609 | 70.61 | 7.10E-94 | 301 | 174 | 27 | 1 | 72 | 245 | 3429489 | 3428983 | 245 | -3 |
| ApnU | JAMABN010000061.1 | GCA_023624855_1_ASM2362485v1_genomic | ApnU | 84.775 | 99.59 | 9.34E-162 | 495 | 289 | 0 | 2 | 1 | 245 | 54354 | 53488 | 245 | -3 |
| ApnU | JAPQKL010000007.1 | GCA_028826915_1_ASM2882691v1_genomic | ApnU | 82.041 | 90.61 | 1.76E-137 | 426 | 245 | 22 | 1 | 23 | 245 | 183921 | 184655 | 245 | 3 |
| ApnU | SGAW01000018.1 | GCA_004153425_1_ASM415342v1_genomic | ApnU | 84.775 | 99.59 | 9.25E-162 | 495 | 289 | 0 | 2 | 1 | 245 | 324004 | 323138 | 245 | -2 |

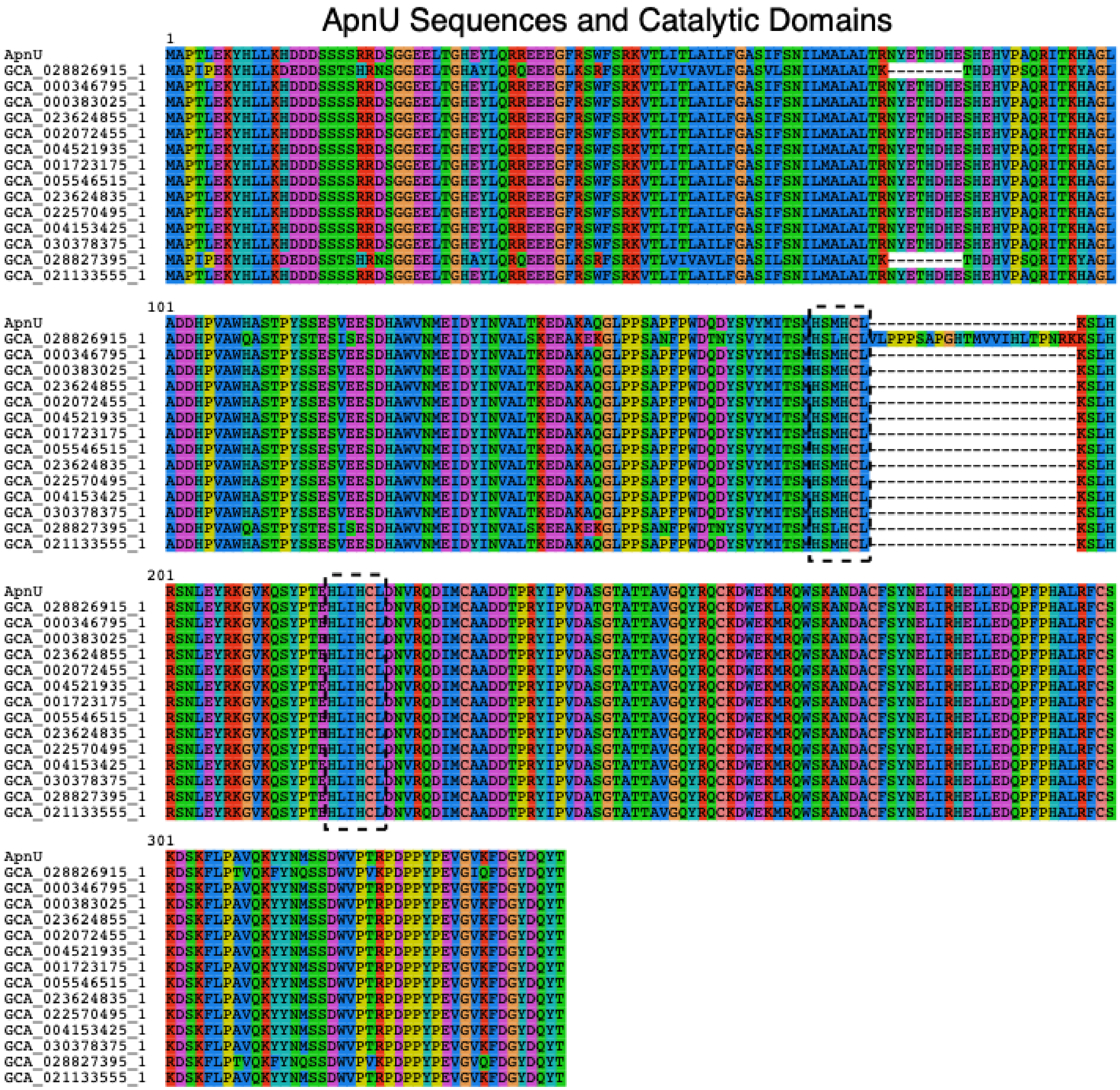

Figure S4. Alignment of the copper-dependent halogenase, ApnU, from the atpenin B biosynthetic gene cluster (MIBiG accession BGC0002067) against putative homologs identified in publicly available *Penicillium* genomes. Key HXXHC motifs are outlined with black dashed lines.

Table S7. SeqForge FASTA-metrics module compared to QUAST assembly metrics output.

| **QUAST** | ***E. coli*** | ***E. coli*** | ***E. coli*** | ***Streptomyces*** | ***Streptomyces*** | ***Streptomyces*** |
|---|---|---|---|---|---|---|
| Assembly | GCA_037575275_1 | GCA_043933725_1 | GCA_047199145_1 | GCA_049606955_1 | GCA_050871035_1 | GCA_050953795_1 |
| # contigs (>= 0 bp) | 8 | 40 | 1 | 5 | 1 | 9 |
| # contigs (>= 1000 bp) | 8 | 11 | 1 | 5 | 1 | 9 |
| # contigs (>= 5000 bp) | 4 | 9 | 1 | 5 | 1 | 8 |
| # contigs (>= 10000 bp) | 3 | 7 | 1 | 4 | 1 | 7 |
| # contigs (>= 25000 bp) | 3 | 7 | 1 | 4 | 1 | 4 |
| # contigs (>= 50000 bp) | 3 | 4 | 1 | 4 | 1 | 3 |
| Total length (>= 0 bp) | 5711995 | 5506486 | 4593097 | 8512991 | 7609140 | 8175220 |
| Total length (>= 1000 bp) | 5711995 | 5493925 | 4593097 | 8512991 | 7609140 | 8175220 |
| Total length (>= 5000 bp) | 5699448 | 5490004 | 4593097 | 8512991 | 7609140 | 8171759 |
| Total length (>= 10000 bp) | 5691234 | 5478789 | 4593097 | 8505710 | 7609140 | 8162927 |
| Total length (>= 25000 bp) | 5691234 | 5478789 | 4593097 | 8505710 | 7609140 | 8116727 |
| Total length (>= 50000 bp) | 5691234 | 5372387 | 4593097 | 8505710 | 7609140 | 8075404 |
| # contigs | 8 | 22 | 1 | 5 | 1 | 9 |
| Largest contig | 5486274 | 5059890 | 4593097 | 6072140 | 7609140 | 7889737 |
| Total length | 5711995 | 5500528 | 4593097 | 8512991 | 7609140 | 8175220 |
| GC (%) | 50.5 | 50.52 | 50.85 | 73.69 | 72.03 | 72.03 |
| N50 | 5486274 | 5059890 | 4593097 | 6072140 | 7609140 | 7889737 |
| N90 | 5486274 | 5059890 | 4593097 | 2010573 | 7609140 | 7889737 |
| auN | 5273174.8 | 4662023.4 | 4593097 | 4820822.2 | 7609140 | 7617090.2 |
| L50 | 1 | 1 | 1 | 1 | 1 | 1 |
| L90 | 1 | 1 | 1 | 2 | 1 | 1 |
| # N's per 100 kbp | 0 | 0 | 0 | 0 | 1.58 | 0 |
| **SeqForge** | ***E. coli*** | ***E. coli*** | ***E. coli*** | ***Streptomyces*** | ***Streptomyces*** | ***Streptomyces*** |
| Filename | GCA_037575275_1 | GCA_043933725_1 | GCA_047199145_1 | GCA_049606955_1 | GCA_050871035_1 | GCA_050953795_1 |
| Num_Contigs | 8 | 22 | 1 | 5 | 1 | 9 |
| Num_Contigs_≥0bp | 8 | 40 | 1 | 5 | 1 | 9 |
| Num_Contigs_≥1kb | 8 | 11 | 1 | 5 | 1 | 9 |
| Num_Contigs_≥10kb | 3 | 7 | 1 | 4 | 1 | 7 |
| Num_Contigs_≥50kb | 3 | 4 | 1 | 4 | 1 | 3 |
| Num_Contigs_≥100kb | 2 | 2 | 1 | 3 | 1 | 2 |
| Total_Length | 5711995 | 5500528 | 4593097 | 8512991 | 7609140 | 8175220 |
| Total_Length_≥1kb | 5711995 | 5493925 | 4593097 | 8512991 | 7609140 | 8175220 |
| Longest_Contig | 5486274 | 5059890 | 4593097 | 6072140 | 7609140 | 7889737 |
| Shortest_Contig | 1902 | 509 | 4593097 | 7281 | 7609140 | 3461 |
| GC_Content(%) | 50.5 | 50.52 | 50.85 | 73.69 | 72.03 | 72.03 |
| N_Count | 0 | 0 | 0 | 0 | 0 | 0 |
| auN | 5273175 | 4662023 | 4593097 | 4820822 | 7609140 | 7617090 |

Table S7 continued.

| **SeqForge** | ***E. coli*** | ***E. coli*** | ***E. coli*** | ***Streptomyces*** | ***Streptomyces*** | ***Streptomyces*** |
|---|---|---|---|---|---|---|
| N50 | 5486274 | 5059890 | 4593097 | 6072140 | 7609140 | 7889737 |
| N90 | 5486274 | 5059890 | 4593097 | 2010573 | 7609140 | 7889737 |
| L50 | 1 | 1 | 1 | 1 | 1 | 1 |
| L90 | 1 | 1 | 1 | 2 | 1 | 1 |

1. Bruna T, Hoff KJ, Lomsadze A, Stanke M, Borodovsky M. Braker2: Automatic eukaryotic genome annotation with genemark-ep+ and augustus supported by a protein database. NAR Genom Bioinform. 2021;3(1):lqaa108.
2. Flynn JM, Hubley R, Goubert C, Rosen J, Clark AG, Feschotte C, et al. Repeatmodeler2 for automated genomic discovery of transposable element families. Proc Natl Acad Sci U S A. 2020;117(17):9451-7.
3. Stanke M, Diekhans M, Baertsch R, Haussler D. Using native and syntenically mapped cdna alignments to improve de novo gene finding. Bioinformatics. 2008;24(5):637-44.
